# A general trait-based model for multiplex ecological networks

**DOI:** 10.1101/2023.08.08.552546

**Authors:** Kayla R. S. Hale, Elisa Thébault, Fernanda S. Valdovinos

## Abstract

Ecological networks can represent the structure of food webs, energy flow, and the many and diverse types of interactions between species in ecosystems. Despite its tremendous importance for understanding biodiversity, stability, ecosystem functioning, research on ecological networks has traditionally been restricted to subsets of the species or interactions in ecosystems, i.e., “subnetworks” such as pollination networks or food webs. As a result, the structure of “multiplex” networks that include multiple interaction types is mostly unknown and there is no robust, underlying theory to support their study. Some ecological traits, such as body size or length of mouth parts, are well-known as key predictors of different species interactions. These traits are often strongly related to each other due to evolutionary history, allometry, and selection, and this relatedness may constrain the structure of ecological multiplex networks. We use this idea to develop a model that simulates multiplex ecological networks by interconnecting subnetworks using correlated traits. Our model predicts how multiplex network structure, measured as the overlaps between species’ functional roles, is affected by neutral processes, interaction rules, and trait constraints, while the structure of individual subnetworks is independent of these trait correlations. Additionally, our model accurately predicts the structure of an observed multiplex network using existing knowledge on species trait correlations and basic features of known ecological subnetworks. This work will stimulate new studies of the structure and dynamics of complex ecosystems by providing a null expectation for how multiplex ecological networks are structured under different ecological conditions.

## Main Text

Ecological networks are a powerful conceptual and technical tool to represent the complexity of species-rich ecosystems. Studying the structure of ecological networks has shown that species interactions are arranged in predictable, nonrandom patterns determined by species’ traits^1–4^; these patterns in turn shape how ecosystems respond to perturbations. For instance, studies^5–10^ show that effects of species extinctions on biodiversity and ecosystem function strongly depend on the traits and position – or functional role – of the removed species in ecological networks, as well as on how species restructure their interactions (“rewire”) under novel conditions. Empirical networks are expensive to document in terms of time and expertise, and their structures are sensitive to the spatial scale, number, and composition of sampled species^11–14^. Stochastic network models have therefore been key for advancing ecological and evolutionary research in complex systems by providing fast generation of simulated replicates that successfully reproduce key features of empirical networks and null expectations to assess the effects of sampling under specific ecological hypotheses^13,15–20^.

However, these models are currently restricted to networks (hereafter, “subnetworks”) composed of only one type of ecological interaction at a time, such as predator-prey or pollination interactions. In contrast, natural systems include diverse types of interactions between co-occurring species, which can be represented as “multiplex” networks of interconnected subnetworks^16,21,22^. The pattern of interconnections between subnetworks, i.e., the functional roles of species that participate in multiple interaction types, can strongly affect ecosystem stability and function^22–25^ by propagating feedbacks between different subnetworks, with, for example, herbivores affecting the structure and dynamics of pollination subnetworks through their impact on plants^26–28^. Studying these feedbacks has been identified as a goal of ecological networks research for more than 15 years^16,19,21,22,24,25,29–31^, but though empirical multiplex networks have begun to be documented^32–38^, the lack of underlying models still limits advancement^22^. Here, we develop a general stochastic network model and illustrate its use for addressing three key objectives in ecological networks research: i) predicting the structure of empirical multiplex networks, ii) understanding the drivers of species’ functional roles across interaction types, and iii) assessing the scaling of multiplex networks with species richness over space.

The multiplex network model we propose builds on “interaction rules” from classic subnetwork models (Fig. 1A-C) by including key “trait correlations” for the species involved in different subnetworks (Fig. 1D-E). Existing models successfully reproduce the structure of empirical subnetworks because, across interaction types, subnetworks are of low dimensionality, meaning that species can be ordered along a single or very few axes to determine who interacts with whom, given a simple set of rules for interaction^1,39^. Our model uses the fact that these abstract axes are well-represented by measurable ecological traits^1,40^, with different traits tending to determine different interaction types (e.g., body size for carnivory, flower and animal mouthpart lengths for pollination, plant chemical compounds for herbivory). Such traits are likely correlated due to various fundamental ecological and evolutionary processes (e.g., genetic correlations, trade-offs in species performances)^27,28,41–45^ and should therefore affect the roles of species in multiple interaction types (see Supplementary Text). For example, the phenolics that attract pollinators and seed dispersers to flowers and fruits also attract herbivores and seed predators, so that the plants who support the most mutualists are also potentially those most vulnerable to consumers^27,28,41^.

**Fig. 1:**
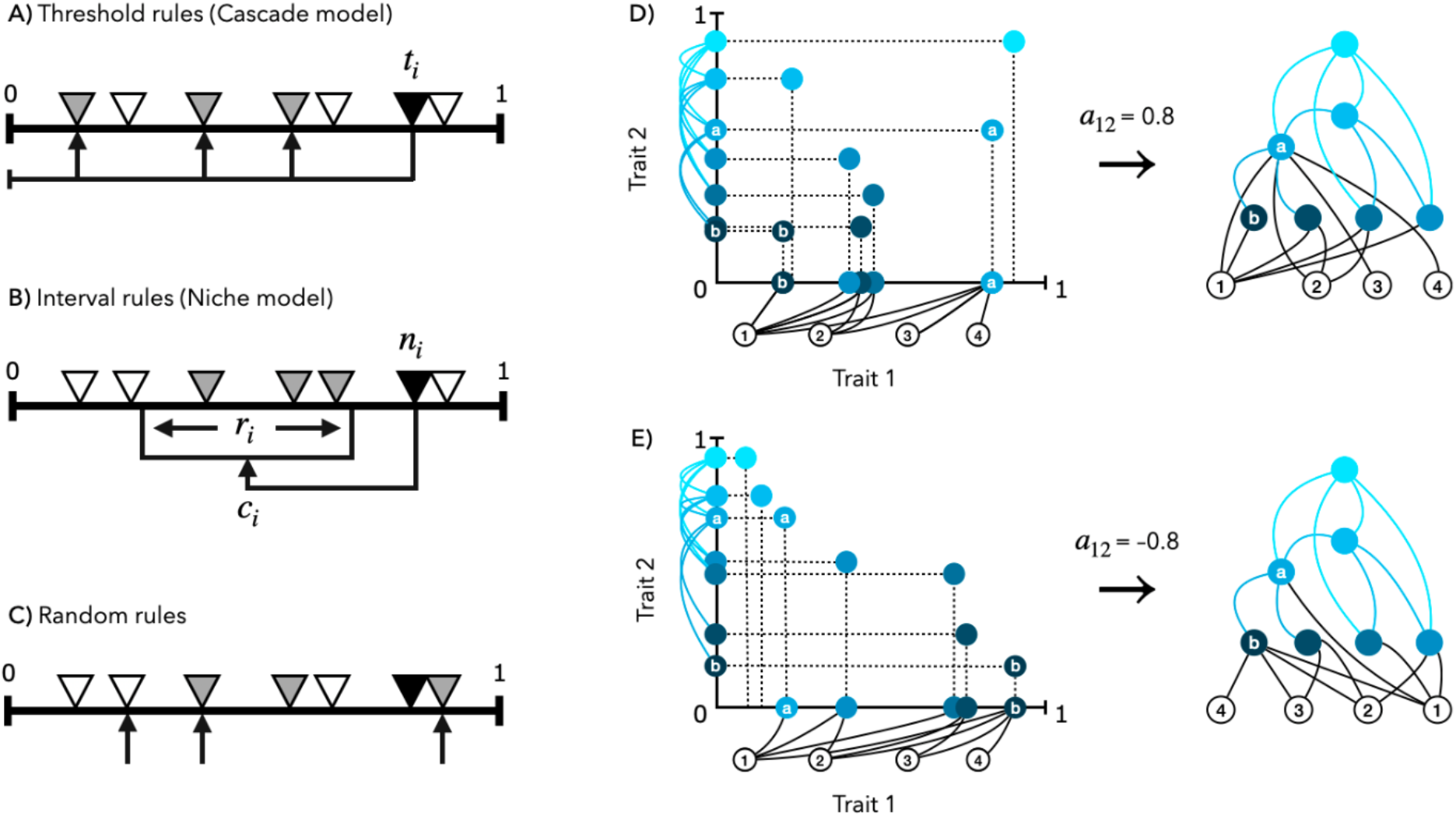
Multiplex networks are constructed from subnetworks with correlated trait axes. (**A-C**) Classic models construct subnetworks by randomly assigning species (triangles) along a single uniform trait axis and linking them (arrows) into unipartite food webs with simple “interaction rules” that embody ecological hypotheses. (**A**) In the Cascade Model^46^, a species (*i*) can potentially interact with any species below its threshold trait value (*t*_*i*_), while (**B**) in the Niche Model^3^, *i* interacts with all species within a range (*r*_*i*_) centered at a trait value (*c*_*i*_) lower than its own (*n*_*i*_). Following^20^, we refer to these models and their extensions to bipartite subnetworks as following “threshold” or “interval rules”, respectively. (**C**) Under the random rule, a species can potentially interact with any species, regardless of trait value. (**D, E**) Toy example of assembling a multiplex network. Trait for *S* = 10 species are randomly sampled from a copula so that traits 1 and 2 are correlated with coefficient *a*_12_ but are distributed uniformly along each axis. Trait 1 determines a bipartite subnetwork where animals (filled nodes) interact with (black links) plants (white nodes) according to threshold rules. Trait 2 determines a unipartite subnetwork where animals interact with each other (blue links) according to interval rules. Horizontal or vertical placement of the nodes indicate their trait values on each axis. Trait correlations do not affect the subnetworks’ emergent structure but rather their interconnection into multiplex networks. For example, when trait axes 1 and 2 have a strong positive correlation (*a*_12_ = +0.8, **D**), high trait-valued species in subnetwork 2 tend to have higher degree (*k*) in subnetwork 1 than low trait-valued species in subnetwork 2 (compare animal *a* with *k*_1_(*a*) = 4 and animal *b* with *k*_1_(*b*) = 1). This is reversed when the trait axes are strongly negatively correlated (*a*_12_ = −0.8, **E**): *k*_1_(*a*) = 1 while *k*_1_(*b*) = 4.

Our model generates a multiplex network of *S* species consisting of *n* subnetworks as follows. For each subnetwork, we define a corresponding trait axis *x* and an “interaction rule” that predicts whether any pair of species interact based on their trait values (see Methods and Fig. 1). Following previous models, the interaction rules we use are: (1) “Threshold” (Fig. 1A), in which a species can potentially interact with any partner below a threshold trait value^20,46^, (2) “Interval” (Fig. 1B), in which a species interacts with all partners within a range (interval) of trait values^3,20^, and (3) “Random” (Fig. 1C), in which species can potentially interact with any partner, irrespective of trait values. These simple rules well-characterize observed ecological interactions^1,2,15,19,20^. For example, pollinators tend to access plants below a threshold of flower corolla length matching the length of the pollinators’ mouth parts (Fig. 1, Trait 1, threshold rule), while predators tend to hunt prey within a contiguous range of body sizes smaller than their own (Fig. 1, Trait 2, interval rule).

Each species *i* ∈ {1, …, *S*} is assigned a set of traits 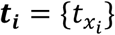 drawn from a “copula,” a multivariate probability distribution in which the values for each trait *t*_*x*_ are uniformly distributed along their axis but constrained by correlations between each pair of traits *a*_*xy*_ (Fig. 1D-E, Methods). For each trait axis *x*, the model randomly samples *S*_*x*_ species and applies the pre-defined interaction rule with connectance *C*_*x*_, the probability of interaction, to generate a subnetwork. The multiplex network emerges from species participating in multiple subnetworks simultaneously according to the underlying correlated trait axes. As the species richnesses of subnetworks *S*_*x*_ increase relative to overall multiplex richness *S*, species become more likely to participate in multiple interaction types simultaneously.

To illustrate our model’s ability to simulate realistic networks, we first study an empirical example of conservation interest. In habitats that are depauperate of other resources, some bird species forage on both nectar and fruits or seeds, acting as both pollinators and seed dispersers (“double mutualists”)^47,48^. Because traits underlying these interactions are positively correlated across all bird species (*a*_*xy*_ ≈ 0.74 for beak widths and lengths)^1,37,49^, we expect that the bird species that are perhaps the most important pollinators in these environments – i.e., generalists, pollinating the most plant species – are doubly important because they are also generalist seed dispersers – dispersing the most plant species. Specifically, given the number of plant and bird species, the connectance of each subnetwork, and the correlation among bird traits, our model reproduces key properties of an empirical pollination and seed dispersal network from the Galapagos Islands^50^ (Fig. 2A). Networks simulated according to threshold rules do not significantly differ from the empirical network in their nestedness, modularity, and variation in birds’ degree (number of plant species) for both the pollination and seed dispersal subnetworks (Fig. 2B), and in their degree distributions for birds both in subnetworks and summed for the full multiplex network (Fig. 2C). Additionally, the expected high overlap between the most generalist pollinators and seed dispersers, as well as the overlaps between other functional roles, closely match the overlaps in the empirical network (Fig. 2D, Extended Data Fig. 1). These results highlight that our model is able to predict the structure of empirical multiplex networks, extending the existing models for subnetwork structure. Furthermore, approaching multiplex networks in this way can operationalize the often interaction-specific knowledge of organisms’ natural histories (e.g., their generality or traits) to predict their simultaneous functional roles in natural ecosystems.

**Fig. 2:**
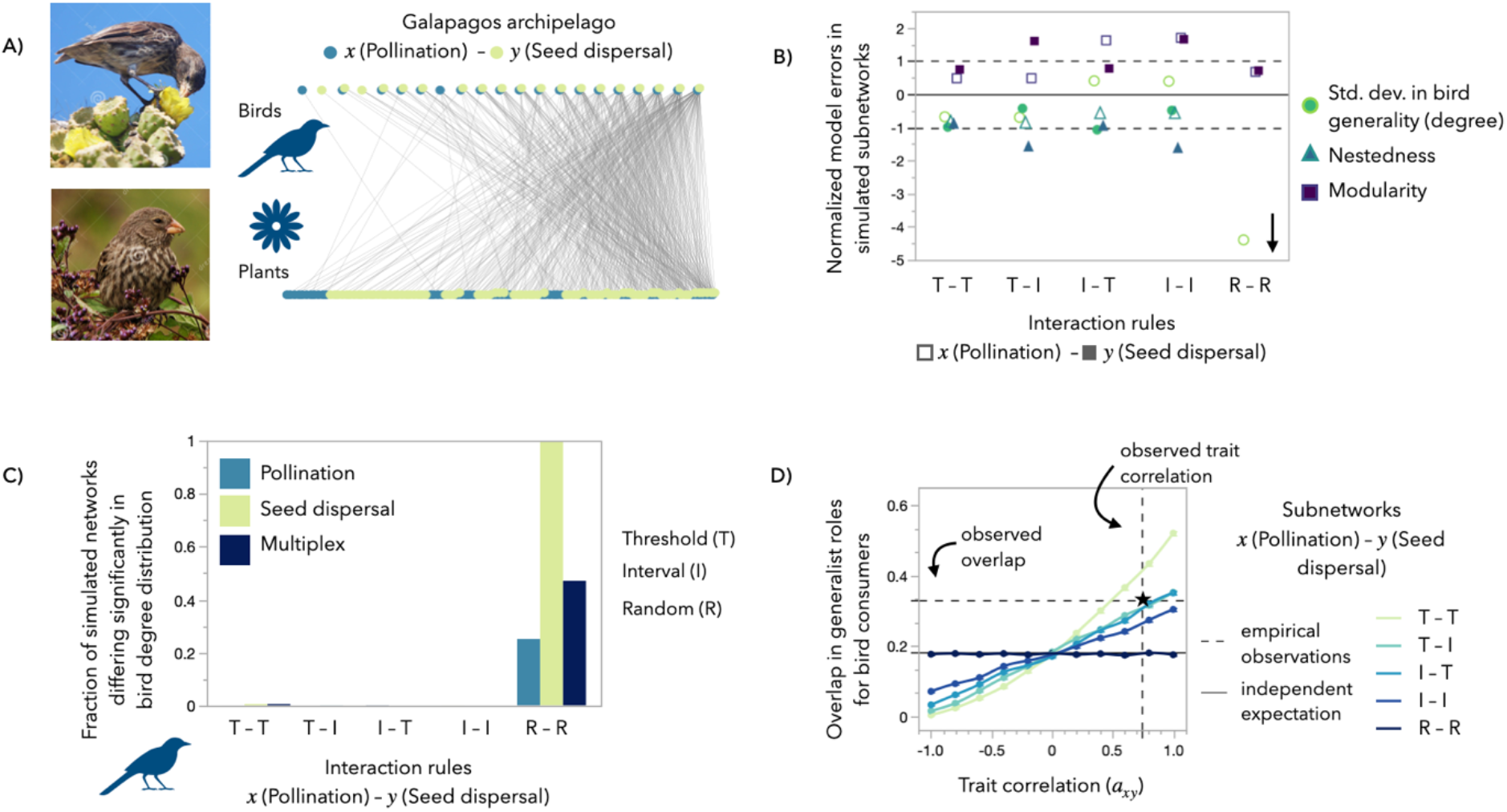
Properties of birds as “double mutualist” seed dispersers and pollinators are well-predicted by the multiplex model with empirical trait correlations. **(A)** An empirical multiplex network of bird-pollination and -seed dispersal interactions from the Galapagos^50^. Offset nodes indicate that the same species engages in both the pollination (blue) and seed dispersal (green) subnetwork. **(B)** Normalized model errors (NMEs) show normalized differences in median nestedness (triangles), modularity (squares) and variability of bird generality (circles) for simulated subnetworks compared to the empirical pollination (hollow) and seed dispersal (filled) subnetworks. NME between (–1, +1) indicates no significant difference at the 95% confidence level. Three points for random rules fall below the y-axis scale (−5 > NMEs > -10, indicated by arrow). **(C)** Fractions of simulated networks for which two-tailed Kolmogorov-Smirnov tests rejected the null hypothesis that degree distributions of simulated and empirical networks were sampled from the same underlying distribution (95% confidence level). **(D)** Empirically-observed overlap (mean Jaccard similarity from 100 bootstrapped replications, dashed horizontal line) between birds’ functional roles as generalist pollinators and seed dispersers compared to overlaps in simulated multiplex networks (colored lines). Generalists are in the top 1/3 of the consumer degree distribution (i.e., linked to the most resource plants). Points (resp. error bars) are means (resp. standard errors) from N = 1,000 multiplex networks simulated for each combination of trait correlation and interaction rules: threshold (T), interval (I), random (R). The black line marks the expected overlap if subnetworks were generated independently (random rules or uncorrelated traits *a*_*xy*_ = 0). The star marks the intersection of the empirical overlap with the empirical correlation between traits thought to underly pollination (beak length) and seed dispersal (beak width) interactions across all bird species^1,37,49^ (*a*_*xy*_ ≈0.74, dashed vertical line). **(B, C)** Calculated for each set of simulated multiplex networks pooled across trait correlations (N = 12,100).

The relationship between species’ functional roles (see Methods) in different subnetworks^27,38,51,52^ or in terms of their participation in modules^21,33,53^ has been central to the study of empirical multiplex networks. While a recent meta-analysis of the few available networks suggests such relationships can vary greatly^38^, our model provides predictions for how overlaps in species’ functional roles across multiple subnetworks depend on trait correlations, interaction rules, and whether species are consumers or resources (Fig. 3, Extended Data Fig. 2). Importantly, these predictions (described below for *n* = 2 subnetworks) are qualitatively robust across ecological complexity (species richness and connectance), including the number of component subnetworks, ≥ 2. At the same time, the structure of subnetworks themselves remain independent of trait correlations.

**Fig. 3:**
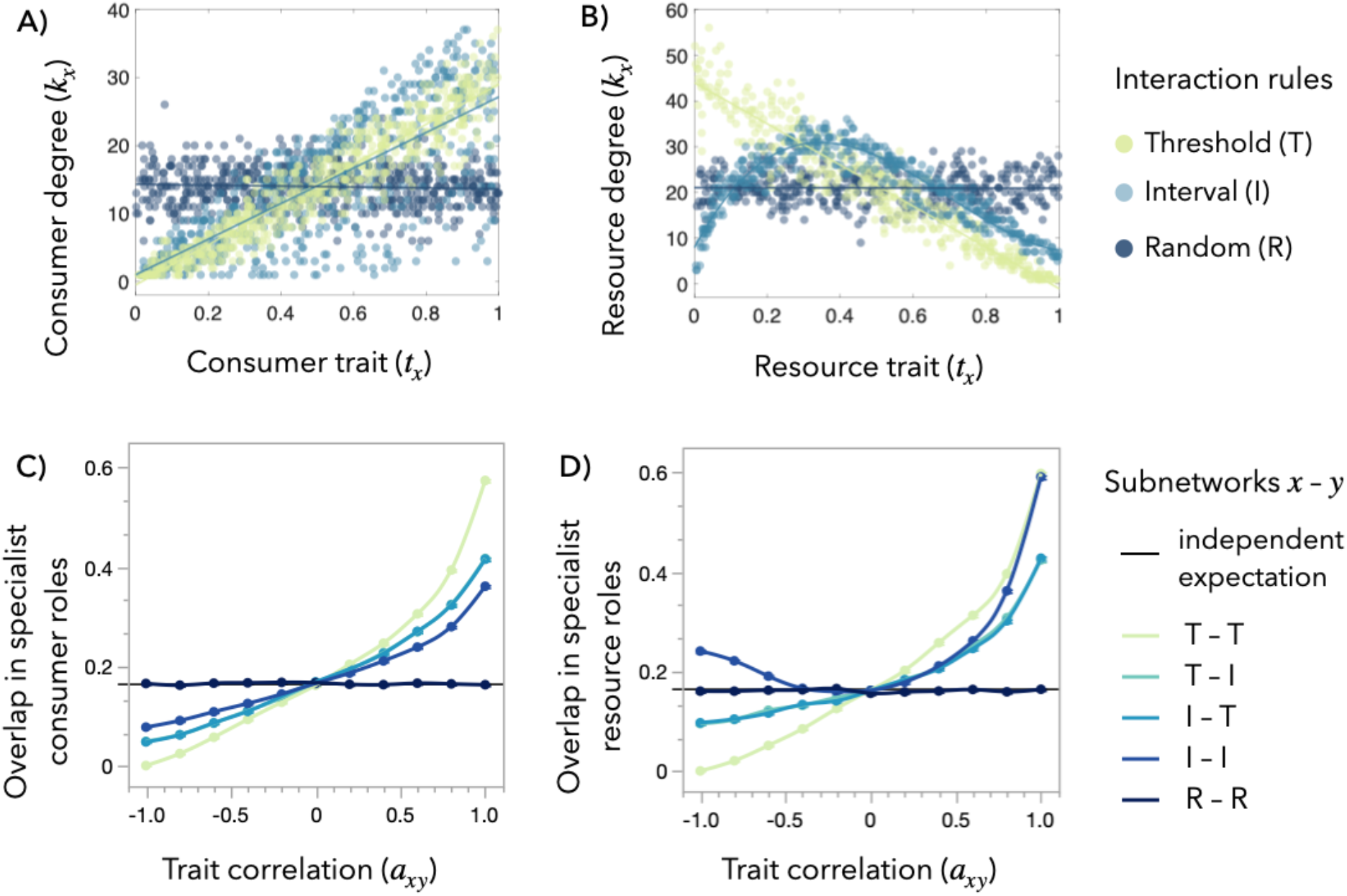
Traits constrain the overlap of species into functional roles. Consumer (**A, C**) and resource (**B, D**) species differ in their patterns of interconnection in multiplex networks. (**A-B**), Interaction rules (Fig. 1) determine the relationship between trait value (*t*) and degree (*k*) in each subnetwork (*x*). These relationships, in turn, determine whether species of a given functional role in subnetwork *x* are the same group of species of a given functional role in subnetwork *y* according to different trait correlations (*a*_*xy*_). That is, the overlap of species into functional roles (calculated using Jaccard similarity, see Methods). (**C-D**) Overlap of low-degree (specialist) consumer (**C**) or resource (**D**) species between subnetworks. Threshold rules result in strong positive or negative relationships between trait and degree for consumer (**A**) or resource (**B**) species, respectively. Interval rules allow both very high and low trait-valued resource species to escape consumption, resulting in lower degree for these resources (**B**). Therefore, low-degree consumers tend to have overlapped functional roles between interval subnetworks when the underlying traits are positively correlated (**C**), but low-degree resources overlap when traits are both positively or negatively correlated (**D**). Overlaps between functional roles follow the analytical expectation (black line) for independently-generated subnetworks when trait correlations are zero or subnetworks are generated according to random rules, regardless of the rules used to generate the other subnetworks (only random-random results shown for clarity). (**A-B**) Points are (**A**) consumer or (**B**) resource (plant or prey) species from N = 10 subnetworks generated using each interaction rules set. (**C-D**) Points (resp. error bars) are means (resp. standard errors) for N = 4,000 multiplex networks composed of *n* = 2 subnetworks, simulated for each combination of interaction rules (colors). See Extended Data Fig. 2 for more combinations of functional overlaps.

When at least one subnetwork is generated by random rules, overlaps in functional roles follow the analytical calculation for overlap among independent sets (Fig. 3C-D). In subnetworks generated by threshold or interval rules, the correlation between species’ degrees tracks the underlying correlation between species’ traits (Extended Data Fig. 3). Accordingly, and as expected from empirical studies ^22,27^, we found high overlap in the same functional roles when the trait axes were positively correlated (*a*_*xy*_ > 0; Fig. 3C-D). That is, specialists in subnetwork (*x*) also tend to be specialists in subnetwork (*y*). An exception to this pattern occurs for resources when subnetworks are generated by the interval rule: counterintuitively, we found positive degree correlations and corresponding high overlaps between low-degree resources at both positive and negative trait correlations (Fig. 3D). That is, the least vulnerable resource species in subnetwork (*x*) also tend to be the least vulnerable in subnetwork (*y*) when traits are positively correlated (*a*_*xy*_ > 0) and strongly negatively correlated (*a*_*xy*_ → −1). Our results also reveal that overlaps in functional roles are not complete (are < 1) – some species will not play the same role in two subnetworks – even when interactions are determined by strongly correlated traits. The overlaps are strongest for threshold rules, weaker for interval rules, and intermediate when one subnetwork is generated by threshold and the other by interval rules. Overlaps are also not commutative: the tendency for, e.g., generalist parasitoids to be specialist pollinators (low degree in *y*) does not necessarily imply the same tendency for generalist pollinators to be specialist parasitoids (low degree in *x*). Together, these results indicate that trait correlations are not always synonymous with overlapping functional roles in multiplex networks, due to the underlying rules and inherent stochasticity of observed ecological interactions. Yet, despite this stochasticity, our results indicate that species that participate in multiple interaction types – the key species that control the propagation of perturbations between subnetworks – have predictable functional roles across subnetworks, given their underlying interaction rules and trait correlations.

An open question in network ecology is understanding how the structure of empirical networks depends on their observational scale, especially the richness of sampled species^12– 14,19,54^. While studies have documented the scaling of subnetwork structure with richness and the consistent rewiring of interactions over time and space^10,14,18,55–57^, such patterns remain unexplored for multiplex networks. Using our model, we studied how neutral sampling of species richness affects local multiplex networks sampled from a regional pool defined by a set of species and traits with an underlying trait structure. We sampled species from this regional pool and applied interaction rules to generate a multiplex network, that is, a local realization of the network assembled from the region (Fig. 4A). Variation among local networks results from turnover in species and rewiring of their interactions. We found that local networks built from ecological interaction rules (i.e., threshold, interval) show less turnover, especially at increasing local network complexity (larger samples from the region), indicating that ecological processes limit the turnover between local networks at the regional scale (Fig. 4B). While realized trait correlations among local species remain relatively independent of local network complexity (Fig. 4C), the overlaps in the species composition of subnetworks increase with local network richness independently from interaction rules and trait correlations (Fig. 4D), as expected from neutral sampling. Additionally, overlaps in species’ functional roles across subnetworks increase with local subnetwork richness while becoming more divergent among interaction rules (Fig. 4E, Extended Data Fig. 4). Together, these results give novel predictions of how richness – through neutral sampling processes – affects observed multiplex network structure. In particular, our prediction of increasing overlap in species composition among subnetworks with richness contradicts the empirical finding^47,48^ that consumers are more frequently involved in multiple subnetworks on depauperate islands (low local richness) than on mainlands (high local richness). This suggests that other ecological and evolutionary processes (e.g., niche filtering, trait evolution) shape such patterns, rather than neutral sampling processes alone.

**Fig. 4:**
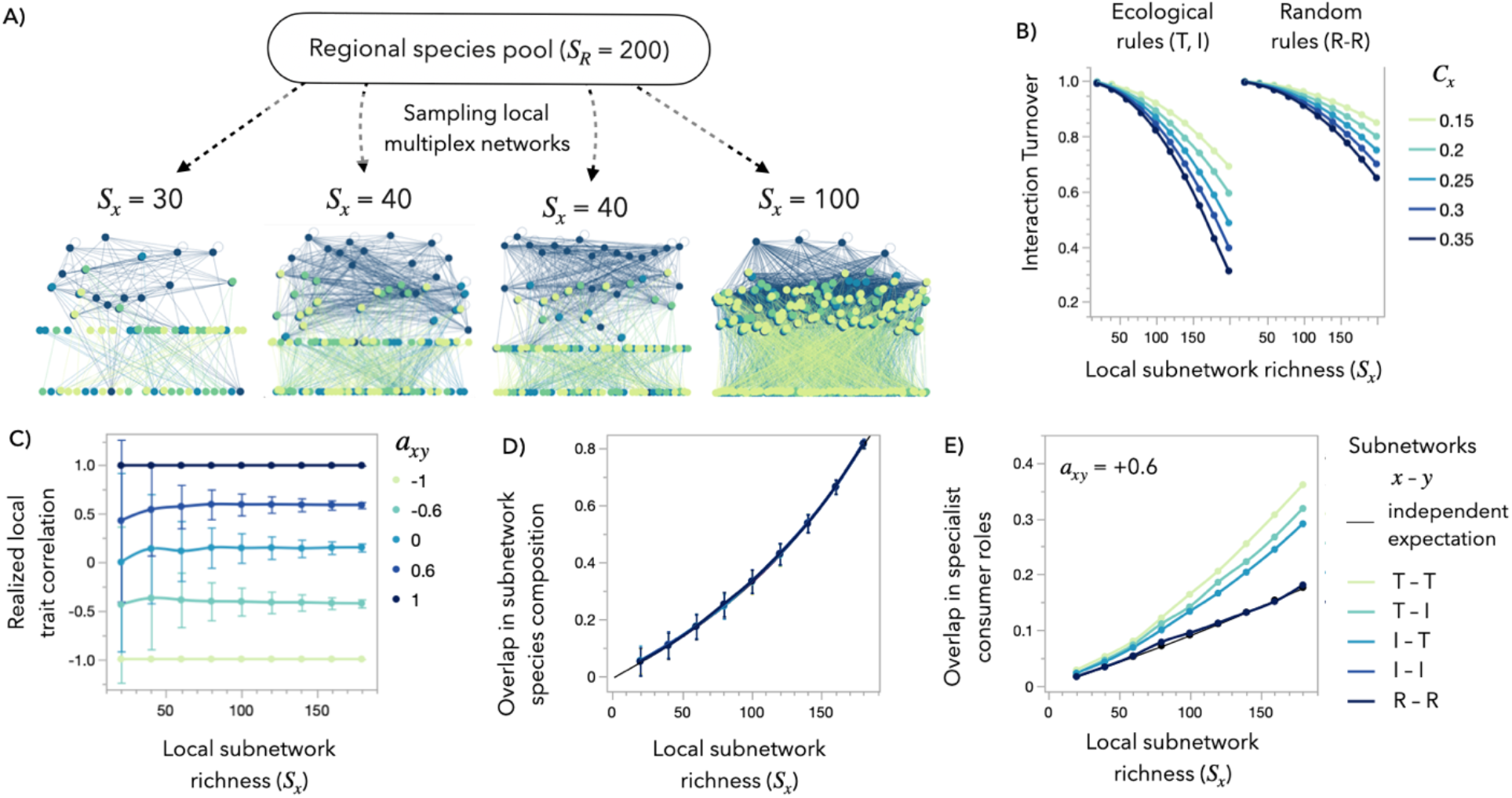
Variation in local multiplex networks assembled from a regional species pool is constrained by ecological interaction rules. (**A**) Multiplex network structure varies due to neutral processes, interaction rules, and trait constraints, corresponding to a wide range of ecological conditions. Our model generates variation from these sources by assembling local multiplex networks from random (neutral) samples of (*S*_*x*_) species into subnetworks from a fixed regional pool (*S*_*R*_). See Methods for parameters of example networks. (**B**) Local networks show decreasing turnover in interactions, calculated as Jaccard dissimilarity, when subnetworks have higher connectance (*C*_*x*_, colors) and contain more of the regional species pool (*S*_*R*_= 200). Multiplex networks generated from ecological interaction rules (threshold [T] or interval [I])) show less interaction turnover than networks generated from random rules. (**C**) The correlation between species’ traits in local networks is representative of the underlying regional trait correlations (*a*_*xy*_, colors), with less variation as local networks approach the regional richness. (**D**) Neutral sampling processes (black line) predict the overlap of species across multiple interaction types, regardless of underlying trait structure or interaction rules (colors). (**E**) Overlaps in functional roles between subnetworks scale with local subnetwork richness, but preserve the qualitative pattern in deviation from neutral expectations (black line for independently-generated subnetworks) given their set of interaction rules (colors). See Extended Data Fig. 4 for more functional overlaps. Underlying data in (**B-E**) is pooled for each treatment from N = 100 multiplex networks of *n* = 2 subnetworks generated according to each set of interaction rules with connectances *C*_*x*_, richnesses *S*_*x*_, and trait correlation *a*_*xy*_. Points are means for (**B**) N = 19,800 (ecological) or 9,900 (random) pairs of networks, (**C**) N = 2,000 networks, (**D**) N = 500 networks, and (**E**) N = 100 networks with *a*_*xy*_ = +0.6. Bars are standard errors in (**B, E**) and standard deviations in (**C-D**).

Our results are dependent upon the suitability of interaction rule sets to simulate the structure of species interactions. We focused on networks of consumer-resource interactions (both trophic and mutualistic) whose generating “rules” have been robustly studied^40^. This may be too restrictive for non-trophic interactions like facilitation and competition that are determined more by spatial proximity than by ecological traits. However, our approach is flexible: any set of interaction rules can be used to generate subnetworks, including extant approaches not studied here^15,58–61^. Indeed, species do not access all resources within a trait range^1,3,59,60,62^, nor can they access all resources under a threshold trait with equal probability, as imposed by the interval or threshold rules, respectively. Real diets likely follow a rule intermediate between these two extremes, which has been the logic behind other generating models for ecological networks (Nested Hierarchy^60^, Minimum Potential Model^59^, Probabilistic Niche Model^62^). Therefore, our results for threshold and interval rules may provide extreme bounds for predictions of multiplex network properties.

Here, we presented a model for multiplex ecological networks that generalizes upon the classic subnetwork models that catalyzed a major body of research on the structure, dynamics, and stability of food webs and other ecological subnetworks^3,16,19,46^. We showed how our model can predict: i) patterns in birds’ functional roles across plant–pollinator and plant–seed-dispersal subnetworks, and the structure of each subnetwork, based on the empirically-known correlation between traits underlying pollination and dispersal interactions in birds, ii) overlaps in species’ functional roles across subnetworks depending on trait correlations, interaction rules, and whether species are consumers or resources; and iii) the structures of local multiplex networks sampled from a broader region. Our model also provides a fast method to generate thousands of multiplex networks across a range of ecological conditions (from the underlying mechanisms of trait constraints, interaction rules, and sampling processes), which may be used to disentangle the seemingly context-dependent results for the relationships between species’ functional roles across different subnetworks^38^ observed in the still-scarce data on multiplex networks currently available. Like classic subnetwork models before, our model can help guide empirical data collection and theoretical study on how different types of species interactions interconnect into multiplex networks and the important ecological and evolutionary consequences of feedbacks between them.

## Methods

### The multiplex network model

#### Species traits

Multiplex networks are a type of “node-aligned multilayer network” in which nodes (species) are potentially present in each layer with perfect coupling and each layer represents a different interaction type^16,21^. Rather than layers, here, we refer to the different interaction types as different “subnetworks” that are interconnected by species that participate in multiple interaction types^22^. We begin by assuming that each of the *n* types of ecological interactions in a community is determined by a single trait (*t*_*x*_). These traits may be measurable or abstract, representing a trait or a composite of multiple ecological traits^1^. Without loss of generality, we assume that each trait can be transformed onto the unit interval [0, 1]. Different traits (*t*_*x*_, *t*_*y*_) may be independent of each other (uncorrelated with rank coefficient *a*_*xy*_ = 0) or constrained (correlated with *a*_*xy*_ ≠ 0) due to, e.g., allometry or evolutionary history. When *a*_*xy*_ = +1 or −1, the traits are perfectly positively or negatively correlated.

To simulate trait values that respect these constraints, we construct a multivariate probability distribution called a “copula” by specifying the correlations between each pair of variables and the overall dependence (the inter-correlation) structure between the variables. The marginal distributions of a copula (i.e., the distributions of each variable in isolation) are uniform on the unit interval. We use either a Gaussian or Student’s *t* dependence structure with 1 degree of freedom to accommodate *n* trait axes simultaneously. The pairwise trait correlations (*a*_*xy*_) parameterize the *n* × *n* trait-axis correlation matrix (***A***); ***A*** must be symmetric, positive semi-definite, and have diagonal entries *a*_*xx*_ = 1 to represent the commutativity of trait correlations. Then, for each species *i*, we can randomly sample the copula, yielding uniformly distributed trait values 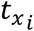 for each of the *n* interaction types that respect our specified correlations. All species have trait values for all the interaction types in the community even if they ultimately do not participate in that interaction.

#### Simulating multiplex networks

In our model, multiplex networks emerge from the interconnection between subnetworks. For each subnetwork (*x*), we specify the species richness for animals (*A*_*x*_) and plants (*P*_*x*_) where *S*_*x*_ = *A*_*x*_ + *P*_*x*_, the number of links between species (*L*_*x*_), and a set of rules for how species’ trait values determine their interactions (see *Interaction rules*, below).

To simulate subnetworks, we randomly sample *S*_*x*_ = *A*_*x*_ + *P*_*x*_ local species from a regional pool of *S*_*R*_ species and apply the selected interaction rules using those species’ trait values. Following classic subnetwork models, we accept the subnetwork if all *S*_*x*_ species have at least one link and the realized *L*_*x*_ is within 3% of the target links. We allow up to 1,000 tries to simulate each subnetwork, sampling *S*_*x*_ species from the regional pool each try.

After successfully simulating all component subnetworks, we check the multiplex network to ensure that every local animal has at least one resource species (i.e., in any subnetwork) if that is appropriate for its ecological interpretation. Note that if all *S*_*x*_ < *S*_*R*_, the unique number of species *S* in the local multiplex network is less than the regional richness *S*_*R*_ because the same species can be sampled into multiple subnetworks. This overlap of species identities into multiple subnetworks causes the interconnection of subnetworks into a multiplex network.

#### Interaction rules

We use interaction rules derived from classic generating models for ecological networks, both in the original unipartite formulations for food webs and in the bipartite versions adapted for mutualistic sub-networks (following^20^). Unipartite: All species in subnetwork *x* can potentially interact with all others so long as their traits are compatible (i.e., meet the criteria of the interaction rule). Subnetwork connectance is 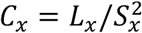. Bipartite: Species in subnetwork *x* are divided into two groups, 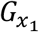 and 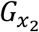. All species in 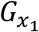 (e.g., animals) can potentially interact with all species in 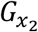 (e.g., plants) so long as their traits are compatible. Subnetwork connectance is 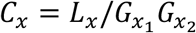, which is *C*_*x*_ = *L*_*x*_/*A*_*x*_*P*_*x*_ when plants and animals are the two groups interacting.

#### Threshold rule

Species *i* with trait value 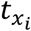 can potentially interact with partner *j* with trait value 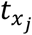 if 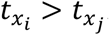. There is a probability *p*_*x*_ that this potential interaction is realized. Unipartite (Original Cascade Model^46^): *p*_*x*_ = 2*C*_*x*_*S*_*x*_/(*S*_*x*_ − 1). Bipartite^20^: Let *N*_*x*_ be the total number of potential interactions in *x*. Then *p*_*x*_ = *L*_*x*_/*N*_*x*_.

#### Interval rule

Species *i* with trait value 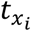 interacts with all partners *j* whose trait 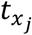 falls within an interval of the trait axis given by 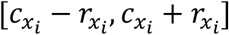, where *i*’s interaction center 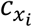 and range 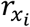 are randomly resampled at each try of simulating a subnetwork. Range 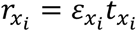, where 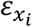 is sampled from a beta distribution with parameters *α*_*x*_ = 1, *β*_*x*_ = (1/2*C*_*x*_) − 1. Center 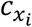 is sampled from a uniform distribution with 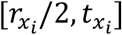. Unipartite (Original Niche Model^3^): All species are assigned interaction ranges and centers. Bipartite^20^: Only species in one group 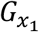 (e.g., animals) are assigned interaction ranges and centers.

#### Random rule^46^

Species *i* can potentially interact with any other species (Unipartite) or with any species in the other group 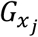 (Bipartite), regardless of trait values. The probability that potential interactions are realized is *p*_*x*_ = *C*_*x*_.

Each of these three rules represents a null hypothesis for subnetwork structures. The threshold and interval rules are ecological hypotheses relating species’ traits to their interactions, while the random rule is the hypothesis that species’ interactions are independent of their trait values.

### Analyzing network structure

#### Multiplex network structure

We develop a new suite of statistics to describe the interconnection of subnetworks via the overlap of species into multiple functional roles. We define these functional roles in a series of categories for each species in the community: presence or absence, animal or plant, consumer or resource, in the top or bottom 1/3 of ranked trait-values, and in the top or bottom 1/3 of ranked degrees within each subnetwork. Degree is the number of links a species has within a subnetwork; it can be separated into incoming-only links, outgoing-only links, or total links. Degree is interpreted as diet breadth, (mutualistic) generality, or vulnerability to consumers, depending on the interaction type and whether the links are incoming or outgoing. We categorize species as having extreme (top or bottom 1/3 ranked) trait values or degree because it is intuitive to consider the overlap between, e.g., the shortest beaked and the most specialist (lowest out-degree) pollinators in a community, and it facilitates the identification of multiplex functional roles.

We use the Jaccard index^63^ (also called the Tanimoto similarity index^64^) to calculate the overlaps between the identity (sets) of species in each functional role both within and between subnetworks:

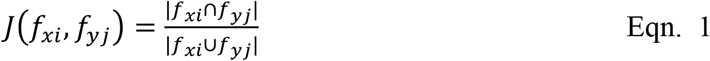

where *f*_*xi*_ is the set of species with functional role *i* in subnetwork *x*. The expected value for the Jaccard index can be calculated analytically as:

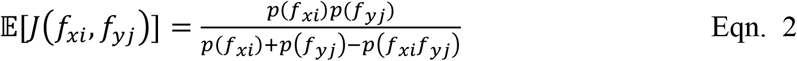

where *p*(*f*_*xi*_) is the probability that any given species is a member of *f*_*xi*_ and *p*(*f*_*xi*_*f*_*yi*_) is the joint probability that any given species is a member of both functional roles. Eqn. 2 reduces to

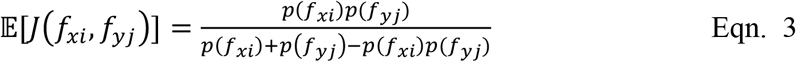

when the functional roles *f*_*xi*_, *f*_*yi*_ are independent^65^, which occurs when the underlying traits are independent (*a*_*xy*_ = 0) or when species’ traits are unrelated to their degree (the random interaction rule). This allows us to establish qualitative expectations in terms of deviation from independence for the functional role overlaps between nonrandom and correlated subnetworks by inspecting their relationships between trait and degree (Fig. 3A-B). Here, we restrict our attention to functional roles in up to 2 subnetworks simultaneously, but a similar approach could be used for *n* > 2 subnetworks.

Because our cutoff of 1/3 for the categorization of species into most extreme trait values or degree is arbitrary, we do not wish to depress our calculation of overlaps between the sets of species in different functional roles only because they contain different numbers of species. Therefore, we calculate overlaps using *m* = min(|*f*_*xi*_|, |*f*_*yi*_|) number of species in each functional role. If multiple species have the cutoff value for being included in functional role, we randomly sample among them if necessary to achieve exactly *m* species total.

To complement previous work^22,27^ we additionally calculate the realized correlation between species’ traits and species’ degrees in each pair of subnetworks, only including species that are present in both subnetworks.

#### Subnetwork structure

We calculate a suite of classic statistics developed for unipartite food webs^15,66^ and bipartite ecological networks^20^ as well as the mean and standard deviation of degree for all species, animal consumers, animal resources (prey), and plants in each subnetwork. We also calculate overlaps between species’ functional roles within subnetworks. For unipartite food webs, we calculate the mean and maximum of short-weighted trophic level (swTL)^67^, the means and standard deviations of herbivory, omnivory, generalism, and vulnerability, the fractions of top, intermediate, and basal species, the predator/prey ratio, the linkage density, the entropy and weighted entropy, and the mean trophic level of the top species only^15,66^. Finally, for bipartite networks, we calculate standardized modularity using the leading eignenvector method^68^ implemented in the BiMat package^69^ and normalized nestedness using the maxnodf package in R^70^.

### Experimental design

#### Analyzing an empirical network

We used our model to analyze an empirical multiplex network (*S* = 129, *A* = 21, *P* = 108) of bird pollination (*S*_*x*_ = 100, *A*_*x*_ = 19, *P*_*x*_ = 47, *C*_*x*_ = 0.148) and seed dispersal (*S*_*y*_ = 100, *A*_*y*_ = 19, *P*_*y*_ = 81, *C*_*y*_ = 0.242) from the Galapagos Islands archipelago. To match the assumptions of the subnetwork models from which we derived the interaction rules, we performed a strict “trophic grouping” on the empirical network so that species with exactly the same links in all subnetworks are grouped into one multiplex “trophic species”. This tended to group specialist plants into trophic species, increasing the connectance (*C*_*x*_ = 0.154, *C*_*y*_ = 0.292) and reducing the richness in the subnetworks (*P*_*x*_ = 44, *P*_*y*_ = 62) and in the overall multiplex network (*S* = 107, *A* = 21, *P* = 86).

We simulated N = 1000 multiplex network replications matched to these richnesses and connectances for the full factorial combination of subnetwork interaction rules {T, I, R} and trait correlations *a*_*xy*_, *p*_*xy*_ ∈ {−1, −0.8, −0.6, …, +1}, where we experimented with using separate underlying trait structures (copulas) for the plants and animals with trait correlations *p*_*xy*_ and *a*_*xy*_, respectively.

Then we analyzed subnetwork and multiplex structure, including overlap of species into functional roles. Plants in this system show less overlap in identity between subnetworks than expected from neutrally sampling the local species pool: empirical overlap *J*(*p*_*x*_, *p*_*y*_)_*emp*_ = 0.233 < expected neutral overlap 𝔼[*J*(*p*_*x*_, *p*_*y*_)] = 0.427 from Eqn. 2, where *p*_*x*_ = *P*_*x*_/*P* = 44/86, *p*_*y*_ = *P*_*y*_/*P* = 62/86. To account for this nonrandom partitioning, we repeated our network simulations with a larger regional plant pool *P*_*R*_ > *P* in order to force the overlap in plant identities between subnetwork to match the empirical value. We calculated the appropriate regional plant richness to be *P*_*R*_ = 136 by solving for *P*_*R*_ in Eqn. 2 with 𝔼[*J*(*p*_*x*_, *p*_*y*_)] = *J*(*p*_*x*_, *p*_*y*_)_*emp*_ = 0.233, *p*_*x*_ = *P*_*x*_/*P*_*R*_ = 44/*P*_*R*_, and *p*_*y*_ = *P*_*y*_/*P*_*R*_ = 62/*P*_*R*_.

To compare empirical and simulated subnetworks, we calculated the standardized modularity, the nestedness, and the mean and standard deviation of degree for all species, animal consumers, animal resources (prey), and plants in each subnetwork. Additionally, we used two-tailed Kolmogorov-Smirnov tests to test the null hypothesis that the degree distribution for bird pollinators, seed dispersers, and summed as double mutualists (multiplex degree) in simulated and empirical networks were sampled from the same underlying distribution.

#### Effect of trait correlations

We simulated multiplex networks across a range of trait-axis correlations and analyzed both the component subnetwork structures and the emergent multiplex network structure.

Specifically, we assembled correlation matrices ***A***_*n* × *n*_ (*n* = 2, 3, … 5) using every combination of *a*_*xy*_ entries {−1, −0.8, −0.6, …, +1} that yields a positive semi-definite matrix. For multiplex networks composed of *n* = 2, 3 subnetworks, we used a full factorial design to simulate N = 1000 replications each for every combination of correlation matrices ***A***_*n*×*n*_ and rules for constructing subnetworks {U, B}×{T, I, R}, where U and B are the unipartite or bipartite application of the threshold (T), interval (I), or random (R) interaction rules. For multiplex networks composed of *n* = {4, 5} subnetworks, we use a reduced design with only 100 replications each for every combination of correlation matrices ***A***_*n*×*n*_ and unique combinations of the threshold (T), interval (I), or random (R) interaction rules where exactly one subnetwork is unipartite (duplicate algorithm combinations arise due to symmetry). As a control, we generated N = 1000 replications of subnetworks (*n* = 1) for all combinations of interaction rules {U, B}×{T, I, R}. For all treatments, we used species richness *S*_*R*_ = *A*_*R*_ + *P*_*R*_ = 120 with *A*_*R*_ = 72 animal species and *P*_*R*_ = 48 plant species for the multiplex network, and *S*_*x*_ = *A*_*x*_ + *P*_*x*_ = 100, with *A*_*x*_ = 60 and *P*_*x*_ = 40 and connectance *C*_*x*_ = 0.35 for each subnetwork *x* = {1, …, *n*}. In Fig. 3, we present only networks with multiplex richness *S* = 116 for clarity, but our results hold for the full range of emergent *S*.

#### Effect of neutral sampling

We generated 5 regional species pool of *S*_*R*_ = *A*_*R*_ + *P*_*R*_ = 200, with *A*_*R*_ = 0.6*S*_*R*_ and *P*_*R*_ = 0.4*S*_*R*_ with underlying trait correlation matrix ***A***_2×2_ with entry *a*_*xy*_ = {−1, −0.6, 0, +0.6, +1}. From each of these regional species pools, we used a full factorial design to generate N = 100 replications of multiplex networks composed of *n* = 2 subnetworks with local richness *S*_*x*_ = *A*_*x*_ + *P*_*x*_ = {20, 40, …, 180} and *A*_*x*_ = 0.6*S*_*x*_, *P*_*x*_ = 0.4*S*_*x*_ species sampled from the regional pool (i.e., with the same trait values in each replication) and assembled according to interaction rules {U, B}×{T, I, R} with connectance *C*_*x*_ = {0.1, 0.15, …, 0.35}.

Variation between multiplex networks simulated according to the same conditions (richness, connectance, traits, and interaction rules) comes from two sources. First, the neutral (random) sampling of species from the regional pool, which allows for differences in species composition between multiplex networks with *S*_*x*_ < *S*_*R*_. Second, from interaction rewiring, which allows the same set of species to potentially engage in different interactions in different observations of the network, as in empirical systems. Interactions are still constrained by species’ traits 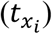, but result from the independent samples of species’ partners in the threshold or random rules, or from species’ range 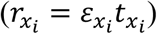 and center 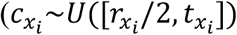 in the interval rules, where ranges 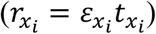 are calculated from the independent sample of 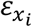 from the same beta distribution.

We quantified interaction turnover (*β*) for each pair of local multiplex networks generated from the same subnetwork richnesses, connectances, and interaction rules and reported the mean for each treatment. Interaction turnover is calculated as Jaccard dissimilarity between the set of links (*l*_*X*_, *l*_*Y*_) in each pair of networks *X, Y*

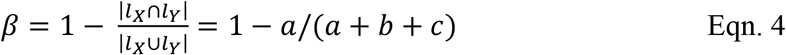

where *a* is the number of shared interactions between networks, and *b* and *c* are the number of unique interactions in each network. This dissimilarity metric includes all sources of interaction turnover from changes in local species composition and rewiring of interactions due to the stochasticity of the interaction rules. Eqn. 4 resembles Eqn. 1 but is its complement (dissimilarity instead of similarity) and quantifies (lack of) overlap in links between multiplex networks rather than overlap in species roles between subnetworks in one multiplex network.

As an illustration of the variation produced by our model, Fig. 4A shows multiplex networks generated from the same regional species pool (*S*_*R*_ = 200, *A*_*R*_ = 0.6*S*_*R*_, *P*_*R*_ = 0.4*S*_*R*_) with differing *S*_*x*_. Each multiplex network is composed of *n* = 4 subnetworks with the following rules: (*x* = 1) bipartite random; (*x* = 2) bipartite threshold; (*x* = 3) bipartite threshold; and (*x* = 4) unipartite interval. All subnetworks were connectance *C*_*x*_ = 0.3, richness *S*_*x*_. From left to right in Fig. 4A, *S*_*x*_ = 30, 40, 40, 120. Subnetworks *x* = 1, 2, 3 were generated with *A*_*x*_ = 0.6*S*_*x*_, *P*_*x*_ = 0.4*S*_*x*_, while subnetwork *x* = 4 was *A*_4_ = *S*_*x*_, *P*_4_ = 0. The parameters of the underlying trait correlations were: *a*_12_ = −0.1, *a*13 = 0.1, *a*14 = 0.1, *a*13 = 0.6, *a*23 = 0.8, *a*34 = 0.6.

## Code availability

Network simulations and topological analyses were performed in MATLAB R2021b (Mathworks) except for nestedness calculations using the maxnodf package in R. Statistical analyses were performed in JMP Pro 16. All code, including sample networks and visualizations, is available online after acceptance.

## End Notes

## Acknowledgements

This work was supported by National Science Foundation grants DEB-2129757 and DEB-2224915 to F.S.V. and the Transatlantic Research Partnership to all authors, a program of FACE Foundation and the French Embassy. This research was supported in part through computational resources and services provided by Advanced Research Computing at the University of Michigan, Ann Arbor. We thank George Kling for comments on the manuscript.

## Author contributions

KRSH developed the model, analysis, and visualization code, and wrote the first draft of the manuscript. All authors contributed to conceiving and conceptualizing this work and revising the manuscript.

## Additional information

Supplementary Information is available for this paper. Correspondence and requests for materials should be addressed to (Kayla R. S. Hale).

## Supplementary Information

### Supplementary Text

Species’ biotic environment – enemies and mutualists – exert selective pressure on multiple traits simultaneously. Organismal allometries and genetic variation limit the traits for a given body plan, while genetic correlations cause evolutionary responses to traits not under direct selection^42,43^. Tradeoffs among traits, such as the competitiveness at exploiting different resources or investments in growth versus reproduction, differentiate species’ niches allowing coexistence^44^. These mechanisms cause correlations between species traits which should also affect the roles of species in multiple interaction types^27,28,45^.

Indeed, we find this to be in the case in the multiplex networks generated by our model under threshold and interval rules. Other rule sets that relate species’ trait values to their ecological role (e.g., generality or centrality in a network) will also show this effect, allowing the prediction of species’ functional roles in multiple subnetworks given some knowledge of the underlying trait constraints. However, trait correlations are tricky: they can indicate a fundamental tradeoff between traits (due either to morphology/physiology or niche partitioning), but they can also depend on the scale of sampling and environmental conditions with meaningful correlations even changing sign across an ecological gradient^42,43,71^. Additionally, species’ traits set the ultimate constraint on their potential interactions and partners, but their observed interactions may be driven more by neutral processes, with e.g., species opportunistically “rewiring” their interactions to the most abundant of their potential partners,

In our model, we find that local trait correlations and qualitative patterns of functional role overlap are preserved across neutral sampling conditions, though deviations from random can become small when regional richness is large compared to the local sample (Fig. 4, Extended Data Fig. 4). In the empirical context, local samples are likely to be a nonrandom subset of the region; rather, they may exhibit traits suitable to the local habitat and, due to observer bias, they are likely the most abundant and least cryptic of the regional species^72^. Thus, we preemptively caution against interpreting model results directly applied to empirical systems. Instead, our model can act as a null expectation for simple ecological hypotheses that guide deeper investigations. Our model could provide further insight into these nonrandom sampling processes by incorporating traits or rule sets^19^ associated with crypticity or abundance onto constrained axes.

## Extended Data Figures & Legends

**Extended Data Fig. 1.**
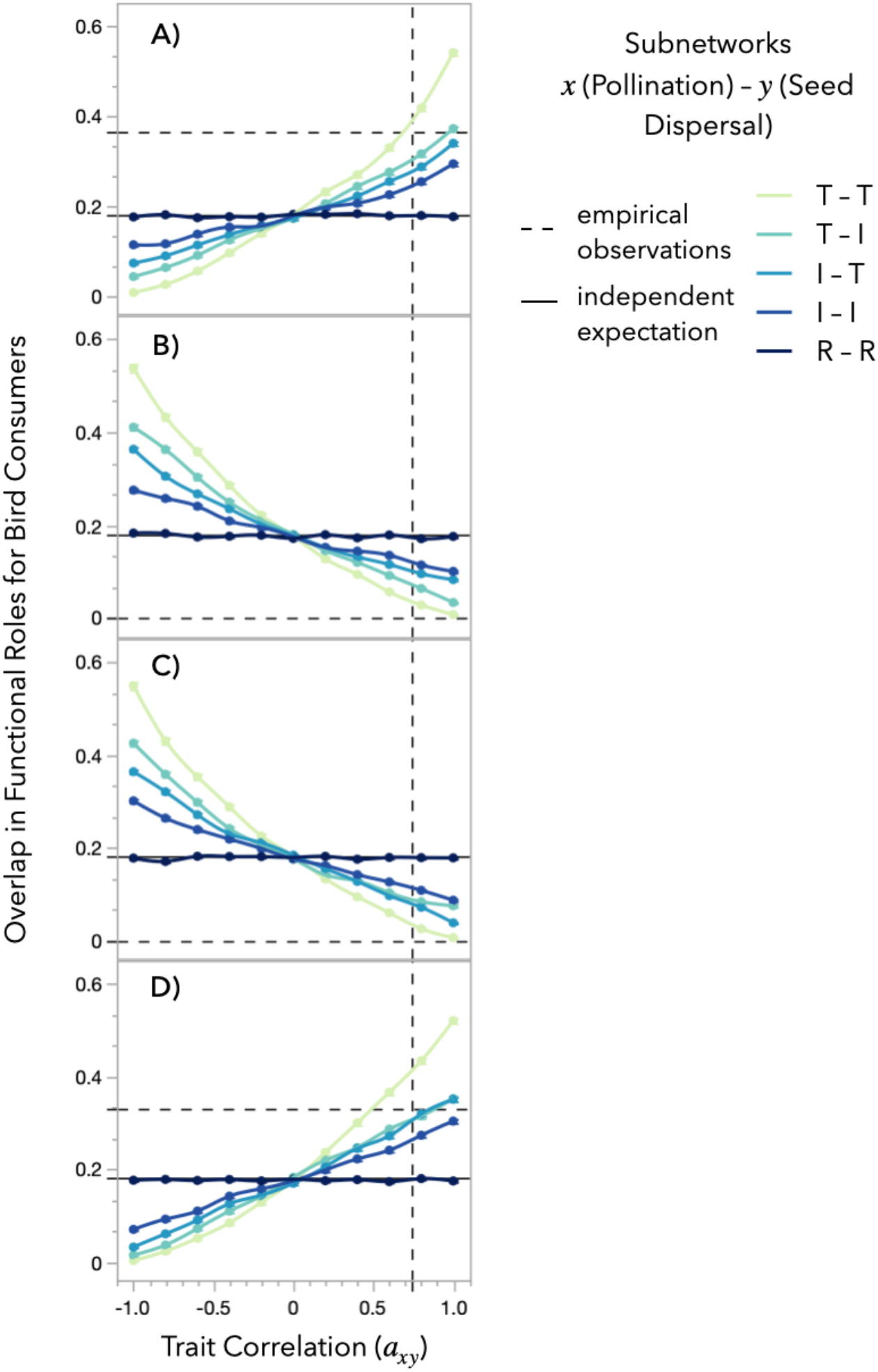
Functional role overlaps for birds in the Galapagos are well-predicted by the multiplex model with nonrandom interaction rules. **(A)** Specialist pollinators–specialist seed dispersers. **(B)** Specialist pollinators–generalist seed dispersers. **(C)** Generalist pollinators–specialist seed dispersers. **(D** Generalist pollinators–generalist seed dispersers. The intersection between the horizontal and vertical dashed lines is the intersection between the observed overlap in the empirical network and the observed trait correlation, respectively. This intersection is the expected value from empirical observations. Points (resp. error bars) are means (resp. standard errors) from N = 1,000 multiplex networks simulated for each combination of trait correlation and interaction rules: threshold (T), interval (I), random (R). The black undashed line marks the expected overlap if subnetworks were generated independently (random rules or uncorrelated traits *a*_*xy*_ = 0).

**Extended Data Fig. 2.**
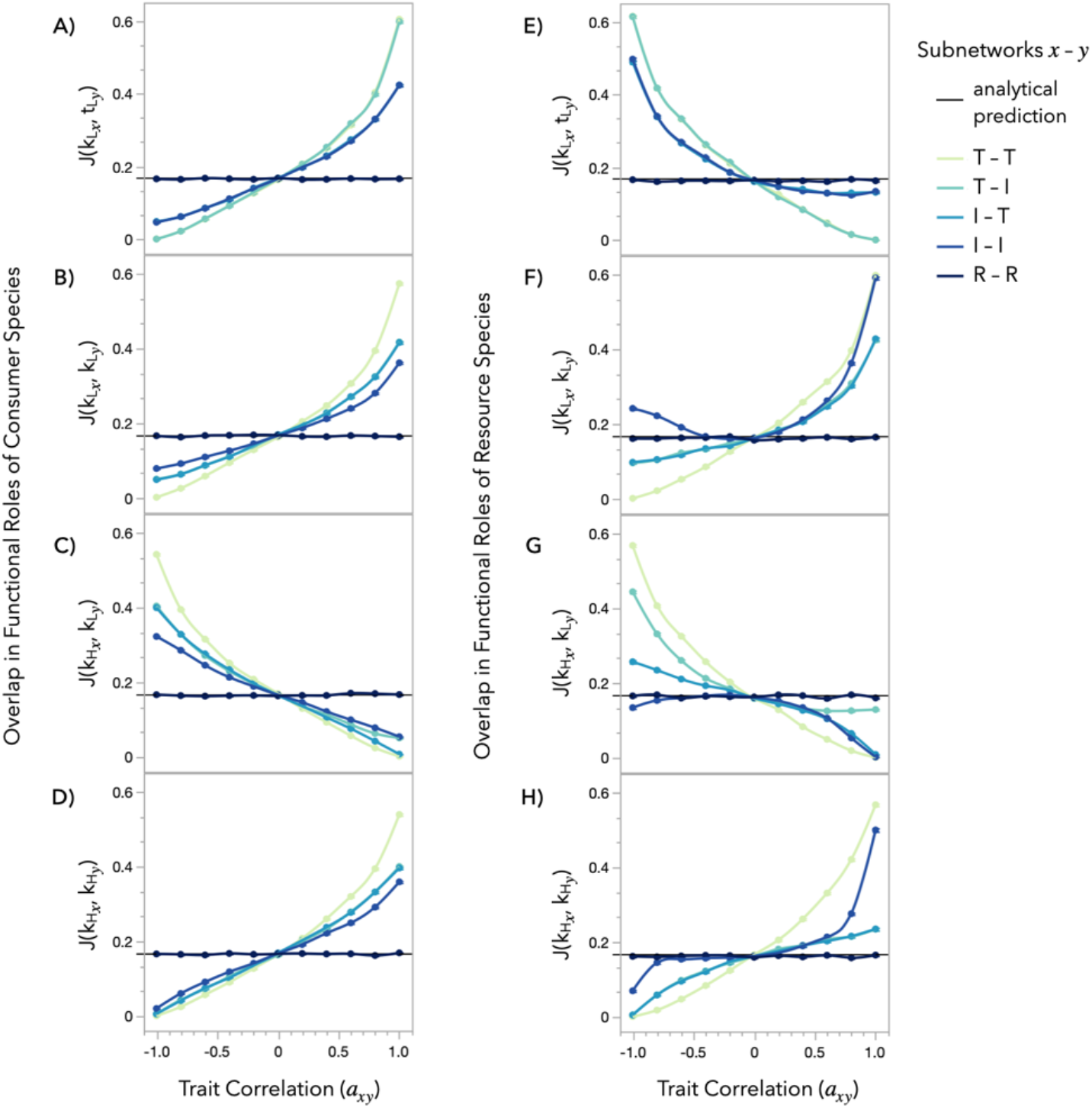
Selected overlaps in functional roles. Interaction rules (Fig. 1) determine the relationship between trait value (*t*) and degree (*k*) in each subnetwork (*x*) for consumers (**A–D**) and resources (**E–H**). These relationships, in turn, determine whether (**A, E**) species with low degree in subnetwork *x* (a functional role in *x*, notated 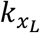) are the same species with low trait value in subnetwork *y* (a functional role in *y*, notated 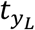) with different trait constraints (*a*_*xy*_). That is, whether there is high overlap of species into functional roles (calculated using Jaccard overlap, *J*, see Methods). (**B, F**) Species overlap between 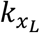 and low degree in subnetwork 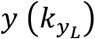 varying with different trait correlations (*a*_*xy*_). Multiplex networks constructed with zero trait correlation or random subnetworks conform to the analytical prediction for the Jaccard overlap (*J*) between independent sets of species (black line). When species interact randomly with respect to their trait values in one subnetwork (ruleset R), the overlaps between species are also independent, regardless of the rules used to generate the other subnetworks (results not shown for clarity). (**C, G**) Species overlap between high degree in subnetwork 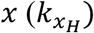 and low degree in subnetwork 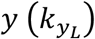 varying with different trait correlations (*a*_*xy*_). (**D, H**) Species overlap between high degree in subnetwork 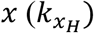 and subnetwork 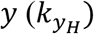. Points (resp. error bars) are means (resp. standard errors) for N = 4,000 multiplex networks composed of *n* = 2 subnetworks, simulated for each combination of interaction rules (colors).

**Extended Data Fig. 3.**
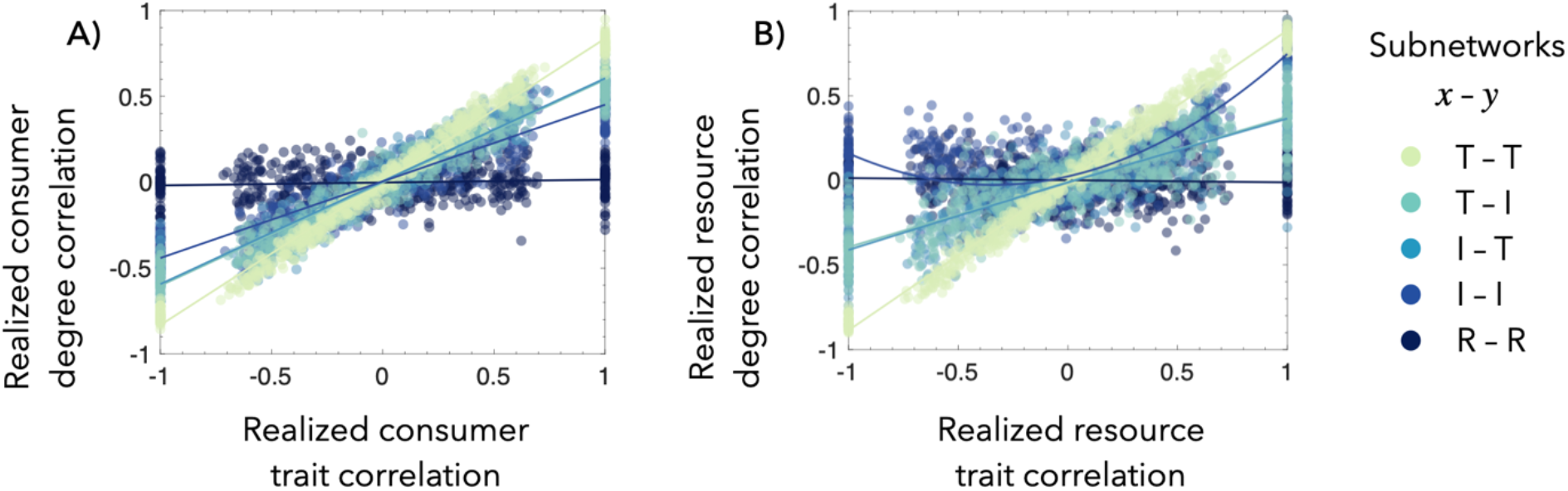
Relationship between trait and degree correlations. The realized correlations among overlapping consumer (**A**) and resource (**B**) species’ degrees track the realized correlations among those same species’ traits for subnetworks generated by ecological (threshold [T] and interval [I]) interaction rules, except that resource species exhibit positive trait degree correlations at negative trait correlations when subnetworks are both generated by interval rules (I–I). Each point (N = 50) is a rank correlation (Kendall’s tau) for the (**A**) consumer or (**B**) resource (plant) species that are present in both of the *n* = 2 bipartite subnetworks in a multiplex network generated according to each interaction rule set (colors).

**Extended Data Fig. 4.**
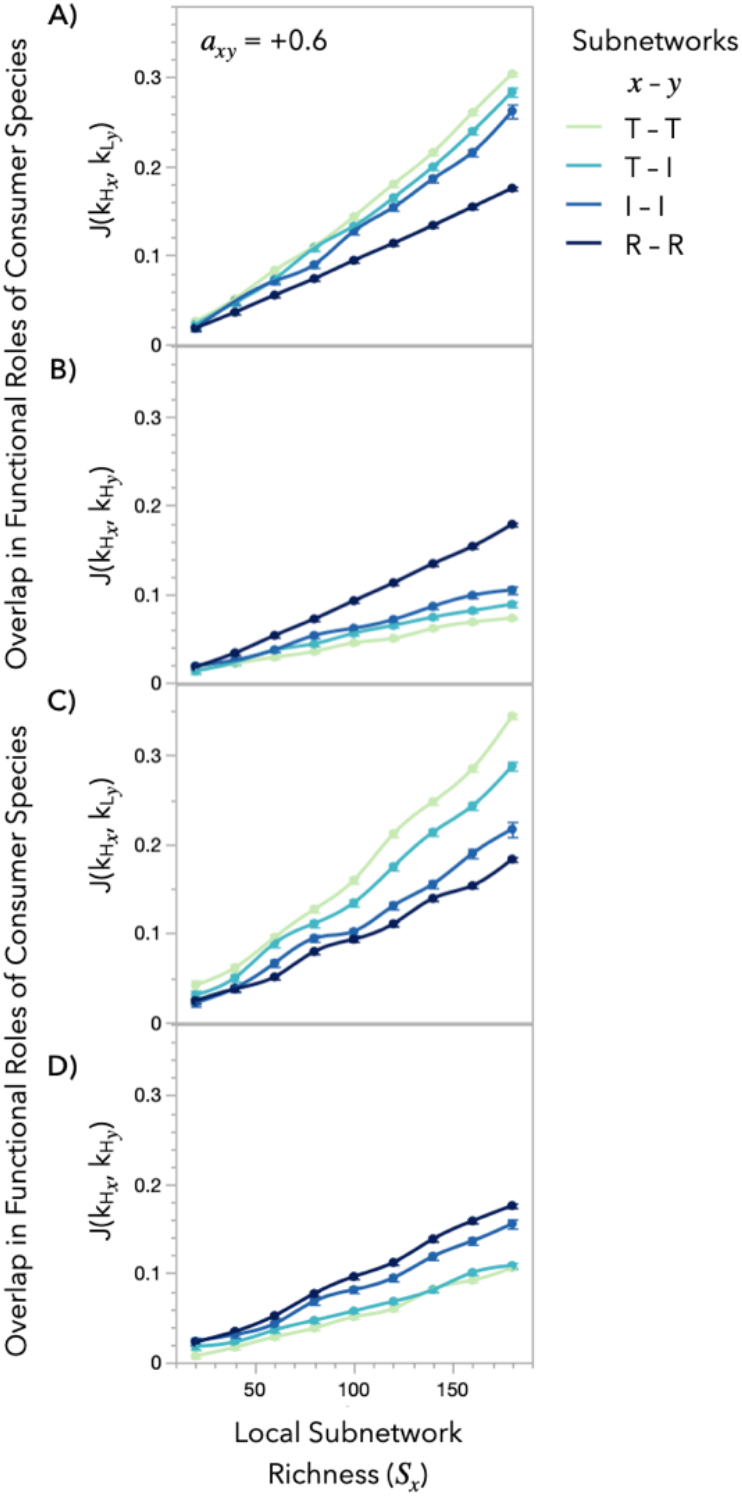
Effect of local richness on functional overlaps. Overlaps in functional roles between subnetworks increase with local subnetwork richness, but preserve the qualitative pattern in deviations between interaction rules (colors). Panels follow the notation from Extended Data Fig 2. Points (resp. error bars) are means (resp. standard errors) for N = 500 multiplex networks of *n* = 2 subnetworks sampled from one underlying regional pool with underlying trait correlation *a*_*xy*_ = 0.6 as an example.

